# Beta Adrenergic Signaling as a Therapeutic Target for Autoimmunity

**DOI:** 10.1101/2025.04.05.647384

**Authors:** Tatlock H. Lauten, Emily C. Reed, Tamara Natour, Lauren J. Pitts, Caroline N. Jojo, Brooke L. Griffin, Adam J. Case

## Abstract

**Background:** We recently identified a molecular mechanism involving beta-adrenergic 1 and 2 receptors (β1/2) in the development of T_H_17 lymphocytes. Pharmacological and genetic inhibition of these receptors in combination, but not separately, impaired the ability of T-lymphocytes to produce proinflammatory interleukin 17A (IL-17A) and instead promoted the production of protective T_reg_ cells that secrete anti-inflammatory interleukin-10 (IL-10). However, it remained unclear whether this regulatory mechanism could serve as a novel therapeutic approach for autoimmune disorders mediated by IL-17A-producing T-lymphocytes.

**Methods:** Multiple sclerosis (MS) is an inflammatory demyelinating disorder of the central nervous system (CNS) characterized by an autoimmune response where both T-lymphocytes and IL-17A are implicated in the pathogenesis of the disease. Using an animal model of MS, termed experimental autoimmune encephalomyelitis (EAE), we addressed the impact of beta adrenergic receptor blockade (genetically and pharmacologically) on EAE disease progression, severity, and T_H_17/T_reg_ balance.

**Results:** The genetic deletion β1/2 receptors, either systemically or specifically in T-lymphocytes, significantly attenuated EAE disease severity and animal weight loss. Pharmacological blockade of β1/2 receptors with either propranolol (lipophilic) or nadolol (aqueous) limited disease severity and weight loss similar to the genetic models. All models showed degrees of shifted T_H_17/T_reg_ balance (suppressing T_H_17 and promoting T_reg_) and decreased T-lymphocyte IL-17A production. Importantly, pharmacological blockade was initiated at the time of symptom development, which mimics the typical time where diagnosis of disease would occur.

**Conclusions:** Our data depict a novel role for β1/2 adrenergic signaling in the control of T_H_17/T_reg_ cells in EAE. These findings provide new insight into the disease progression as well as provide a potential new pharmacological therapy for IL-17A-related autoimmune diseases.

## INTRODUCTION

The connectivity between the central nervous and immune systems is undeniable. Examples of this are highlighted by recent discoveries demonstrating that neurons both release and respond to cytokines, while immune cells do the same with neurotransmitters (1). Additionally, early observations of neuroimmune communication revealed that lymphoid organs appear to be innervated primarily by catecholaminergic nerve fibers (2–5), supporting the idea of neuronal-immune regulation. Indeed, evidence over the last several decades has provided insight into the critical role of the autonomic nervous system in regulating immune cell activity, particularly of T-lymphocytes.

One of the most well-characterized neuroimmune pathways is known as the cholinergic anti-inflammatory reflex (6–11). In this unique pathway, systemic inflammation triggers splenic nerve activation via connections with the vagal nerve. Counterintuitively, while the vagal nerve is parasympathetic in nature, the splenic nerve is exclusively catecholaminergic, thus this vagal response is transmitted via release of sympathetic catecholamine neurotransmitters such as norepinephrine (NE) to the spleen. The released NE binds beta adrenergic receptors on specific subsets of T-lymphocytes causing them to produce and release acetylcholine, which ultimately suppresses inflammation from nearby proinflammatory innate immune cells. However, this model is not without controversy with others suggesting that the vagus nerve is not involved in the process or that the anti-inflammatory effects are due to direct catecholamine binding to immune cells (12–14).

Regarding the latter point, there are many reports of sympathetic neurotransmitters directly impacting inflammatory immune cells, specifically T-lymphocytes. Early studies demonstrated that splenic nerve terminals are in tight contact with T-lymphocytes, and form a type of neural-immune synapse that suggests direct regulation of these cells via neural communication (4). Work over the last few decades has produced a breadth of reports of catecholamine regulation of T-lymphocytes (1, 2). Despite this, no consensus regulatory pattern exists regarding how catecholamines regulate T-lymphocytes due to their responses being highly dependent upon contexts such as timing, location, dosage, and activation/polarization state of the cells.

Nonetheless, we have recently uncovered and reported a regulatory pattern of T-lymphocytes via catecholamine signaling that does appear to be potentially conserved (15). In this work, we identified that genetic or pharmacological inhibition of both beta 1 and 2 adrenergic receptors (β1/2; together, but not individually) significantly decreased interleukin 17A (IL-17A)-producing T-lymphocytes (T_H_17), while increasing forkhead box P3 (Foxp3)-positive T-lymphocytes (Treg) and interleukin 10 (IL-10) production both ex vivo and in vivo. The activation of β1/2 receptors led to the downstream production of cyclic AMP (cAMP), activation of protein kinase A (PKA), and ultimately activation of the cyclic AMP response element-binding protein (CREB) transcription factor; the latter having been also shown to be essential in T-lymphocyte IL-17A generation (16). As proof of principle, we further demonstrated that the inhibition of β1/2 signaling on T-lymphocytes was sufficient to suppress IL-17A production (while promoting IL-10) in a model of psychological trauma (15), suggesting previously unexplored therapeutic implications of β1/2 adrenergic antagonism in IL-17A-related diseases.

IL-17A has been strongly linked to the pathogenesis of various autoimmune diseases including psoriasis, rheumatoid arthritis, and multiple sclerosis (MS) (17–19). IL-17A has a well-defined role in MS, which is a neurodegenerative autoimmune disorder where the myelin coating of neurons is aberrantly targeted by the immune system. T-lymphocyte produced IL-17A orchestrates the disruption of the blood brain barrier and recruitment of innate and adaptive immune cells that ultimately lead to the central demyelination in this disease (20, 21). Moreover, the use of IL-17A neutralizing antibodies has shown encouraging data in both preclinical models of MS as well as in preliminary human trials (22, 23). Therefore, novel therapeutic strategies that limit IL-17A, such as β1/2 antagonism, may have clinical utility in autoimmune diseases like MS.

To explore this further, we investigated the impact of β1/2 adrenergic antagonism on the development of experimental autoimmune encephalomyelitis (EAE); a preclinical mouse model of MS. Utilizing three different methods of β1/2 blockade (i.e., whole body β1/2 knock-outs, adoptively transferred β1/2 knock-outs, and β1/2 pharmacological antagonism), we observed significant attenuation of EAE disease progression. These observations were accompanied by a significant attenuation of IL-17A and T_H_17 cells with a concurrent increase in Treg cells. However, not all models demonstrated the same phenomena to similar extents, suggesting the potential for complex dynamics of β1/2 antagonism. Overall, these data highlight a previously undescribed and critical role for β-adrenergic signaling in the T_H_17/Treg balance in a preclinical model of MS, and provide a novel therapeutic approach (direct or adjuvant) for IL-17A-related autoimmune diseases.

## MATERIALS AND METHODS

### Mice

Wild-type C57BL/6J (#000664; shorthand WT), β1/2 adrenergic receptor knock-out (#003810; shorthand β1/2^-/-^), and Rag2 knock-out mice (#008449; shorthand Rag2^-/-^) were obtained from Jackson Laboratories (Bar Harbor, ME, USA). All mice were bred in house to eliminate shipping stress and microbiome shifts, as well as co-housed with their littermates (≤5 mice per cage) prior to the start of experimentation to eliminate social isolation stress. Mice were housed with standard pine chip bedding, paper nesting material, and given access to standard chow (Teklad rodent diet #8604, Inotiv, West Lafayette, IN, USA) and water ad libitum. Female experimental mice between the ages of 9-13 weeks were utilized in all experiments as outlined by EAE protocols. Male mice were excluded from investigation due to their reduced susceptibility to EAE. Experimental mice were randomized, and when possible, experimenters were blinded to the respective cohorts until the completion of the study. Mice were sacrificed by pentobarbital overdose (150mg/kg, Fatal Plus, Vortech Pharmaceuticals, Dearborn, MI, USA) administered intraperitoneally. All mice were sacrificed between 7:00am and 9:00am Central Time to eliminate circadian rhythm effects on T-lymphocyte function. All procedures were reviewed and approved by Texas A&M University Institutional Animal Care and Use Committee.

### Adoptive transfer

Wild-type or β1/2^-/-^ mice spleens were harvested and T-lymphocytes from spleens were isolated by negative selection, as previously described (24). Briefly, splenocytes in a single cell suspension were negatively selected for T-lymphocytes using the EasyStep Mouse T-Cell Isolation Kit, (StemCell Technologies #19851, Vancouver, BC, USA). Cell viability was measured via a Bio-Rad TC20 Automated Cell Counter using trypan blue exclusion (average yield is approximately 5×10^6^ cells/mouse and purity achieved is >90%). For the adoptive transfer experiments, 5×10^6^ T-lymphocytes were injected intraperitoneally into each recipient Rag2^-/-^ mouse and allowed two weeks for T-lymphocyte engraftment prior to EAE induction.

### Experimental autoimmune encephalomyelitis (EAE) induction

The MOG35-55/CFA Emulsion PTX kit (#EK-2110 Hooke Labs, Lawrence, MA, USA) was used to induce EAE, according to the manufacturer’s protocol. Mice were immunized subcutaneously at two sites on their back with myelin oligodendrocyte glycoprotein 35-55 (MOG35-55) emulsified in complete Freund’s adjuvant (CFA). Subsequently, pertussis toxin (PTX) was injected intraperitoneally twice: 2 and 24 hours after the prior immunization with MOG35-55/CFA at a dose of 100 ng/mL. EAE-induced mice were randomly allocated to each treatment group. The EAE clinical scores and weights were recorded daily as follows: 0, no obvious signs of disease; 0.5, partially limp tail/tip of tail is limp; 1, completely limp tail; 1.5, limp tail and hind leg inhibition; 2, limp tail and complete paralysis of one hind limb or wobbly walk; 2.5, limp tail, dragging hind legs with paralysis of one limb and partial paralysis of the other hind limb; 3, limp tail and complete paralysis of both hind limbs; 3.5, limp tail, complete paralysis of both hind limbs, and can’t right itself from side position; 4, limp tail, complete paralysis of both hind limbs and partial paralysis of one front limb; 4.5, complete paralysis, mouse is not alert and/or able to move around; and 5, death.

### In vivo treatment regimens

β1/2 receptor antagonists propranolol (#P8688, Sigma Aldrich, St. Louis, MO, USA) and nadolol (#HY-B0804, MedChem Express, Monmouth Junction, NJ, USA) were reconstituted in sterile isotonic (0.9%) saline solution and delivered by subcutaneous osmotic minipumps (ALZET Osmotic Pumps, #1002-100uL, Cupertino, CA, USA) using a previously established therapeutic dose of 10 mg/kg (25). Drug or control (i.e., saline only) minipumps were surgically implanted subcutaneously in mice seven days following EAE immunization.

### T-lymphocyte Immunophenotyping

Flow cytometric immunophenotyping was performed as previously described (15, 26). Briefly, spleens and inguinal lymph nodes were harvested, physically dissociated into single cell suspensions, and erythrocyte depleted. Cells were incubated at 37°C for 4 hours in RPMI media supplemented with phorbol 12-myristate-13-acetate (PMA; 10 ng/mL), ionomycin (0.5 mg/mL), and BD GolgiPlug Protein Transport Inhibitor (containing brefeldin A; 1 mg/mL; BD Biosciences, Franklin Lakes, NJ, USA). Following this, cells were washed and resuspended in PBS containing an amine-reactive viability dye (Live/Dead Fixable Cell Stain, #L34960, Thermo Fisher Scientific, Houston, TX, USA) for 30 minutes at 4°C. Antibodies targeting extracellular markers were then added prior to fixation/permeabilization (#00-5523-00, eBioscience, San Diego, CA, USA) and addition of antibodies targeting intracellular markers. Specific antibodies utilized: CD3ε PE-Cy7 (#25-0031-82, Thermo Fisher Scientific, Houston, TX, USA), CD8 Alexa Fluor 488 (#69-0041-82, Thermo Fisher Scientific, Houston, TX, USA), CD4 eFluor 506 (#69004182, Thermo Fisher Scientific, Houston, TX, USA), IL-17A BV711 (#407-7177-82, Thermo Fisher Scientific, Houston, TX, USA), Foxp3 PE-Cyanine5.5 (#35-5773-82, Thermo Fisher Scientific, Houston, TX, USA), IFNy APC-eFluor780 (#47-7311-82, Thermo Fisher Scientific, Houston, TX, USA), and IL-4 BV421 (#562915, BD Bioscinces, Franklin Lakes, NJ, USA), LIVE/DEAD Fixable Far Red (#L34974, Thermo Fisher Scientific, Houston, TX, USA). Cells were assessed using a 4-laser Attune NxT flow cytometer (Thermo Fisher Scientific, Houston, TX, USA), while data was analyzed using FlowJo analysis software (BD Biosciences, Franklin Lakes, NJ, USA).

### T-lymphocyte recall assessment

Briefly, spleens and inguinal lymph nodes were harvested from EAE mice, physically dissociated into single cell suspensions, and erythrocyte depleted. Cells were seeded at 2.0×10^5^ cells per well in a 96-well plate and incubated at 37°C for 72 hours in RPMI media (10% Fetal Bovine Serum, 10 mM HEPES, 2 mM Glutamax, 100 U/mL penicillin/streptomycin, and 50 μM of 2-mercaptoethanol) supplemented with MOG35-55 (10 ug/mL; #DS-0111, Hooke Labs, Lawrence, MA, USA) to induce in vitro recall responses of T-lymphocytes isolated from MOG35-55/CFA-immunized mice. Cell counts and viability were measured via a Bio-Rad TC20 Automated Cell Counter using trypan blue exclusion. Media was harvested for extracellular cytokine analysis and normalized to viable cell counts.

### Cytokine Analysis

Cytokines in plasma and media were measured as previously described (15, 26). Briefly, cytokine levels were assessed using custom Mesoscale Discovery U-Plex kits (Mesoscale Discovery, Rockville, MD, USA) per manufacturer’s instructions. U-Plex kits assessed cytokine levels of IFNy, IL-10, IL-17A, IL-1B, IL-2, IL-21, IL-22, IL-4, IL-6, and TNFa. Analysis was performed using a Mesoscale QuickPlex SQ 120 and analyzed using Mesoscale Discovery software.

### Statistical Analysis

A total of 117 animals were utilized for the studies described herein. The breakdown of these numbers is as follows: 79 WT, 25 β1/2^-/-^, 13 Rag2^-/-^. All data are presented as mean ± standard error of the mean (SEM) with sample numbers displayed as individual data points and N values are included within figure legends where appropriate. Data were analyzed using Mann-Whitney U-test, Wilcoxon signed-rank test, 1-way ANOVA, and 2-way ANOVA where appropriate (specific tests identified in the figure legend). All statistics were calculated using GraphPad Prism (V10, GraphPad).

## RESULTS

### T-lymphocyte β1/2 adrenergic receptors mediate EAE severity

To investigate the effects of beta adrenergic signaling on EAE disease progression and severity, we utilized three main paradigms. The utilization of whole body β1/2^-/-^ mice, adoptive transfer of splenic T-lymphocytes from WT and β1/2^-/-^ donors into Rag2^-/-^ immunodeficient hosts, and antagonism of β1/2 adrenergic receptors via propranolol or nadolol. First, EAE induced in whole body β1/2^-/-^ animals demonstrated slower progression and improved final disease scores over the course of EAE development. Additionally, whole body β1/2^-/-^ animals maintained a more consistent weight throughout the disease progression compared to WT animals (**Figure 1A-B**).

**Figure 1.**
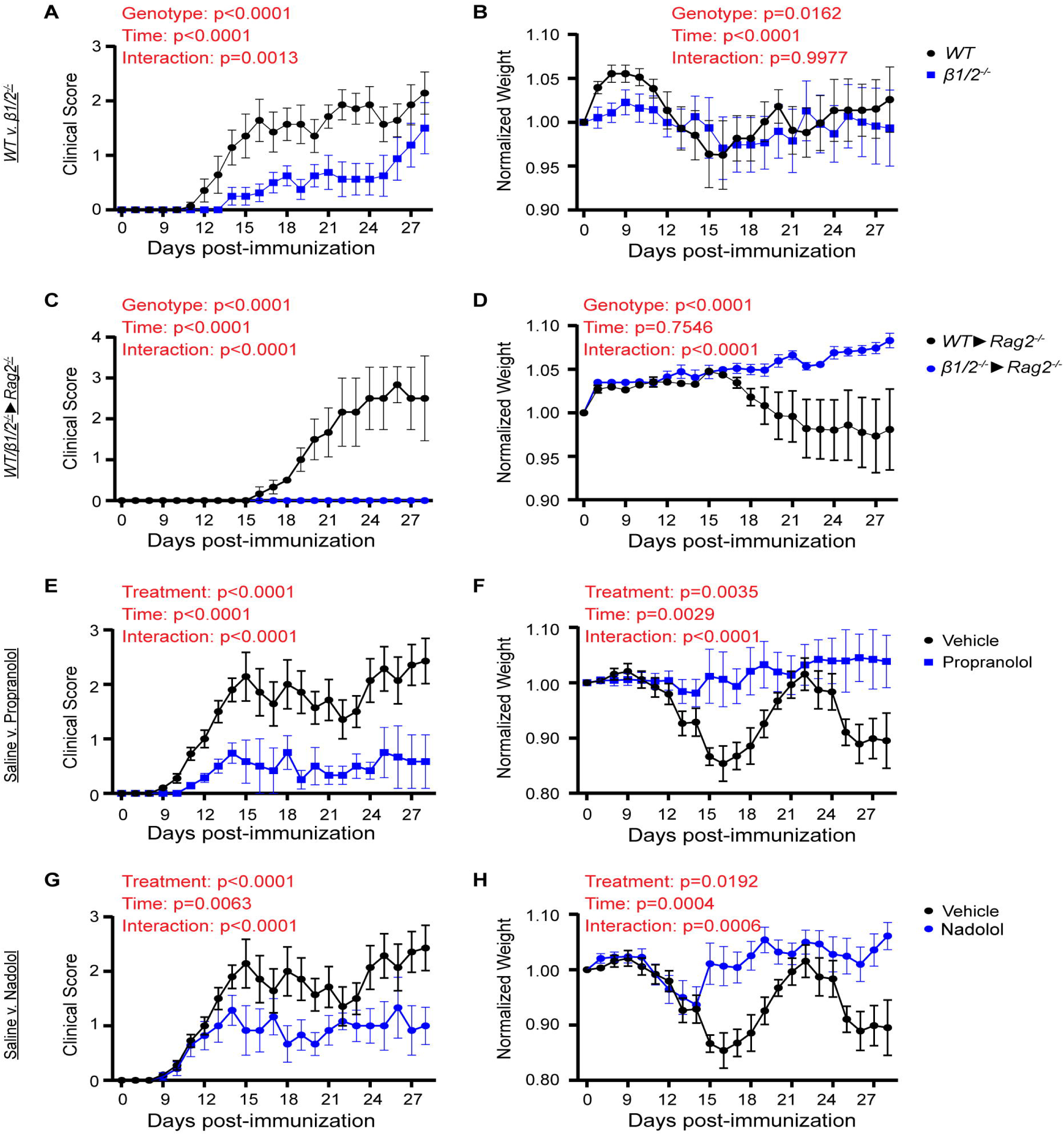
β1/2 adrenergic blockade attenuates clinical EAE score and EAE-induced weight loss. EAE was induced in WT, β1/2^-/-^, or adoptively transferred Rag2^-/-^ mice and were clinical scored and weighed daily. **(A-B)** Whole body β1/2^-/-^ animals versus WT (n=31). **(C-D)** Adoptive transfer of β1/2^-/-^ T-lymphocytes vs WT T-lymphocytes into immunodeficient Rag2^-/-^ mice (n=13). **(E-F)** Pharmacological adrenergic blockade using propranolol (lipophilic β1/2 antagonist) (n=41) or **(G-H)** nadolol (aqueous β1/2 antagonist) (n=34). Statistics by 2-way ANOVA with Bonferroni post-hoc.

To eliminate possible β-adrenergic effects from non-T-lymphocyte cells during EAE, we next utilized T-lymphocyte-deficient Rag2^-/-^ host animals and adoptively transferred splenic T-lymphocytes from WT and β1/2^-/-^ donor animals. Rag2^-/-^ mice receiving WT T-lymphocytes demonstrated the characteristic increase in EAE disease progression and severe weight loss after induction. However, in Rag2^-/-^ hosts that received β1/2^-/-^ T-lymphocytes, disease onset and progression was completely abolished. Surprisingly, Rag2^-/-^ hosts receiving β1/2^-/-^ T-lymphocytes gained weight over the course of disease while the Rag2^-/-^ hosts receiving WT T-lymphocytes lost a significant amount of weight with increasing disease progression (**Figure 1C-D**).

Last, to investigate the efficacy of β1/2 adrenergic blockade as a potential therapeutic in preventing EAE disease progression and severity, β1/2 antagonists, propranolol and nadolol, were utilized. Animals given propranolol one week after EAE induction exhibited markedly reduced development and progression of EAE disease compared to vehicle-treated controls, with some animals exhibiting little to no evidence of disease development **(Figure 1E)**. Additionally, the propranolol-treated animals retained their weight throughout the course of disease as compared to vehicle treated controls who lost weight in a cyclical nature as disease became increasingly severe **(Figure 1F)**. Nadolol treated animals exhibited a smaller impact on EAE progression compared with propranolol in reducing disease progression. However, nadolol treated animals exhibited less severe EAE progression than vehicle treated controls (**Figure 1G**). Nadolol also worked effectively in decreasing weight loss, whereas animals slightly gained weight while on nadolol as compared to saline controls (**Figure 1H)**.

### β1/2 adrenergic receptors on T-lymphocytes regulate balance of T_H_17/T_reg_ during EAE

We next assessed the T_H_17/T_reg_ balance during EAE disease progression in both spleen and lymph nodes across our three main paradigms. Counterintuitively, whole body β1/2^-/-^ animals had increased T_H_17 and decreased T_reg_ cells in the spleens compared to the WT animals **(Figure 2A-B)**. However, the inguinal lymph nodes from β1/2^-/-^ animals possessed the opposite: decreased T_H_17 and increased T_reg_ cells compared to WT (**Figure 2C-D)**. In the adoptive transfer model, both spleens and lymph nodes from Rag2^-/-^ hosts receiving β1/2^-/-^ T-lymphocytes demonstrated decreased T_H_17 cells, but increased T_reg_ cells were only observed in the spleens (**Figure 2E-H**). Last, propranolol was again shown superior to nadolol by significantly decreasing T_H_17 and increasing T_reg_ cells in both the spleens and lymph nodes of animals, whereas nadolol was only able to accomplish this feat in the spleens (**Figure 2I-L**). Together, these data support that T-lymphocyte β1/2 adrenergic blockade alters the T_H_17/T_reg_ balance and correlates with EAE disease severity.

**Figure 2.**
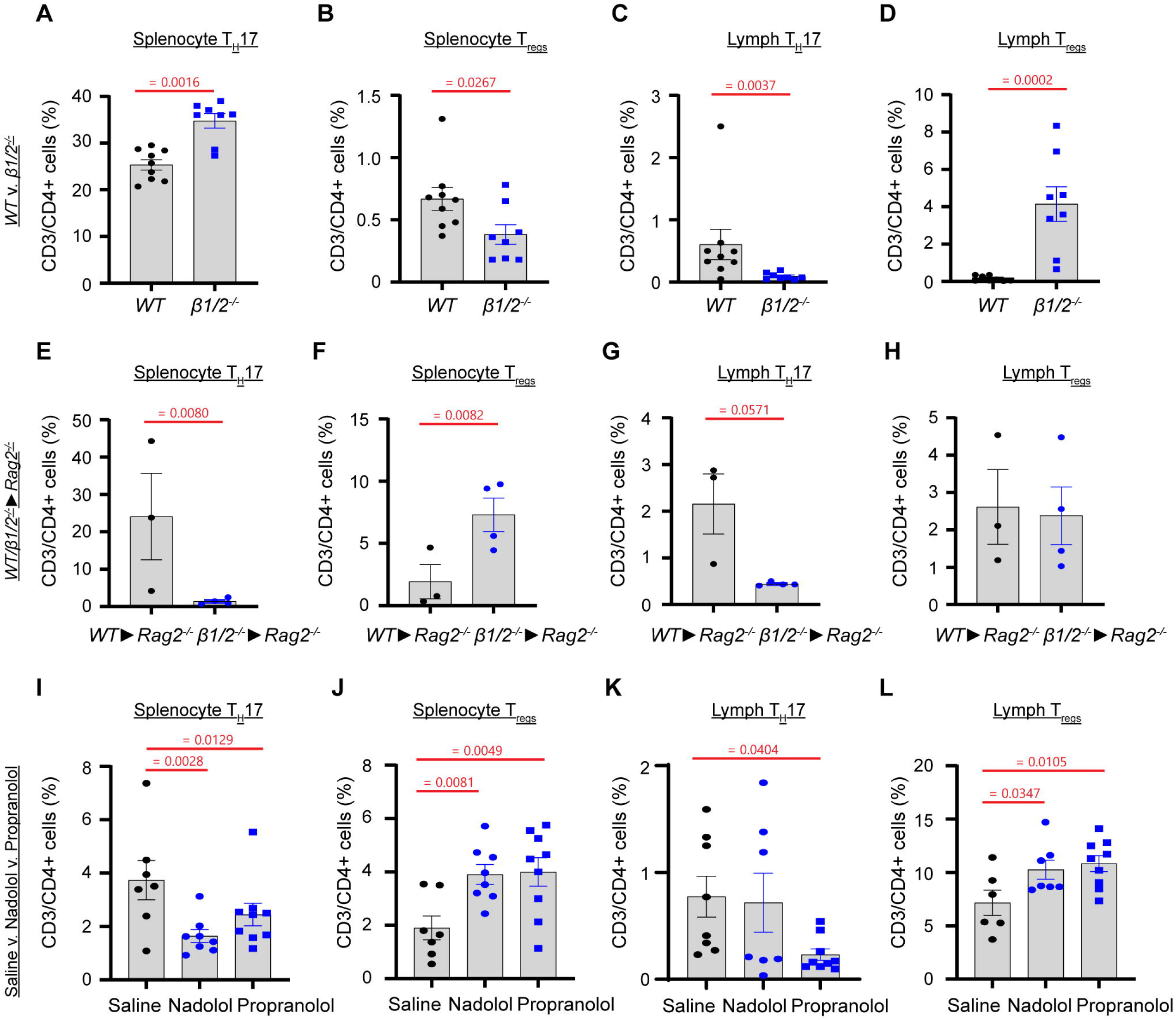
β1/2 signaling alters the T_H_17/T_reg_ balance in EAE. EAE was induced in WT, β1/2^-/-^, or adoptively transferred Rag2^-/-^ mice. Mice were sacrificed at maximal disease severity (day 14-21 depending on model), and spleens and inguinal lymph nodes harvested for T-lymphocyte immunophenotyping. Whole body β1/2^-/-^ animals versus WT splenic T_H_17 (**A**), splenic T_regs_ (**B**), lymph node T_H_17 (**C**), and lymph node T_regs_ (**D**). Adoptive transfer of β1/2^-/-^ T-lymphocytes vs WT T-lymphocytes into immunodeficient Rag2^-/-^ mice splenic T_H_17 (**E**), splenic T_regs_ (**F**), lymph node T_H_17 (**G**), and lymph node T_regs_ (**H**). Pharmacological adrenergic blockade using propranolol (lipophilic β1/2 antagonist) or nadolol (aqueous β1/2 antagonist) splenic T_H_17 (I), splenic T_regs_ (**J**), lymph node T_H_17 (**K**), and lymph node T_regs_ (**L**). Statistics by Student’s t-test (**A-H**) and 1-way ANOVA with Dunnett’s post-hoc (**I-L**).

### β1/2 adrenergic receptors impact T-lymphocyte IL-17A production in EAE

Circulating levels of IL-17A varied across the three paradigms utilized in these investigations. Both whole body β1/2^-/-^ knock-out mice as well as Rag2^-/-^ mice receiving β1/2^-/-^ T-lymphocytes showed significantly decreased plasma levels of IL-17A compared to their respective controls (**Figure 3A, D**). In contrast, both propranolol and nadolol treated animals did not show decreases in circulating IL-17A compared to vehicle controls (**Figure 3G**). This latter observation is currently unexplained, especially given that both these antagonists were able to lower systemic levels of IL-17A previously in a mouse model of psychological trauma (15).

**Figure 3.**
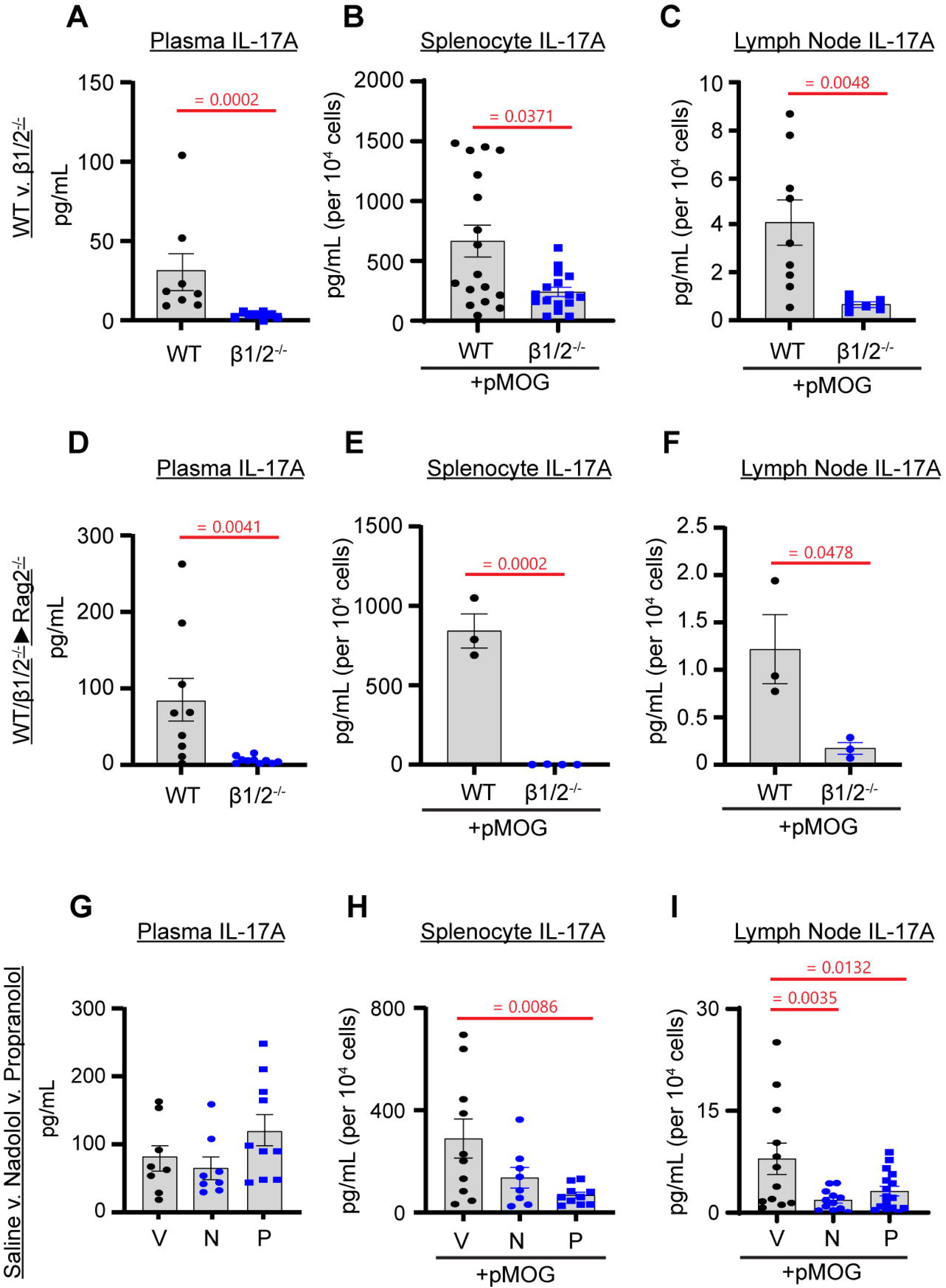
β1/2 adrenergic blockade inhibits T-lymphocyte IL-17A. EAE was induced in WT, β1/2^-/-^, or adoptively transferred Rag2^-/-^ mice. Mice were sacrificed at maximal disease severity (day 14-21 depending on model). Plasma was harvested for circulating cytokine analysis. Spleens and inguinal lymph nodes harvested for T-lymphocyte recall with 10 µg/mL of MOG peptide for 72 hours. Whole body β1/2^-/-^ animals versus WT plasma IL-17A (**A**), splenocyte recall (**B**), lymph node recall (**C**). Adoptive transfer of β1/2^-/-^ T-lymphocytes vs WT T-lymphocytes into immunodeficient Rag2^-/-^ mice plasma IL-17A (**D**), splenocyte recall (**E**), lymph node recall (**F**). Pharmacological adrenergic blockade using propranolol (lipophilic β1/2 antagonist) or nadolol (aqueous β1/2 antagonist) plasma IL-17A (**G**), splenocyte recall (**H**), lymph node recall (**I**). Statistics by Student’s t-test (**A-F**) and 1-way ANOVA with Dunnett’s post-hoc (**G-I)**.

To understand how beta adrenergic signaling impacts the T-lymphocyte’s ability to recognize a secondary immune challenge, we performed antigen recall on total cells from both the spleen and lymph nodes 14-days post EAE induction in all three paradigms. Comparable with their plasma levels of IL-17A, both whole body β1/2^-/-^ knock-out mice as well as Rag2^-/-^ mice receiving β1/2^-/-^ T-lymphocytes demonstrated decreased ability to produce IL-17A from either spleen or lymph nodes upon antigen recall (**Figure 3B-C, E-F**). Interestingly, propranolol showed a similar decrease in IL-17A production in both spleens and lymph nodes, whereas nadolol only significantly impacted the lymph nodes (**Figure 3H-I**). These results suggest that pharmacological beta antagonists may have preferential sites of action on T-lymphocytes involved in EAE progression without impacting circulating levels systemically. Taken together, these data in whole put forth strong evidence that β1/2-adrenergic receptors alter T-lymphocyte-mediated IL-17A production, and hold promise as a potential new therapeutic target in IL-17A-releated autoimmune diseases.

## DISCUSSION

The interaction between the autonomic nervous system and immune regulation has garnered increasing attention in recent years, particularly regarding T-lymphocyte function. Norepinephrine (NE), a key neurotransmitter in the sympathetic branch of the autonomic nervous system, plays a critical role in modulating immune responses through its interaction with adrenergic receptors, especially β-adrenergic receptors on T-lymphocytes. Among these, β2-adrenergic receptors have been identified as primary mediators of NE’s effects on T-lymphocyte activity, influencing cytokine production and differentiation (1, 27). However, growing evidence suggests that other adrenergic receptors, including β1 and even alpha, may also contribute to T-lymphocyte function, particularly in the context of inflammatory immune responses (28, 29).

The β2 receptor is the most prominently studied adrenergic receptor on T-lymphocytes given its high expression on these cells (30). Early work using β2 agonists on naïve cells overwhelmingly demonstrated suppression of IL-2 and other proinflammatory cytokines (31), which led to the prevailing concept that sympathetic signaling was immunosuppressive regarding T-lymphocytes. While this may be true in reductionist settings, the dynamic nature of T-lymphocyte activation, polarization, interaction with other cell types, as well as the expression of adrenergic receptors makes the rule of immunosuppression by the sympathetic nervous system not as universal as it was once perceived. Moreover, many of these studies were performed during a time when only IFNγ and TNFα were investigated from T-lymphocytes, which overlooked the diverse repertoire of potential cytokines regulated by the sympathetic nervous system. Additionally, reports of T-lymphocyte control by other adrenergic receptors further add to the complexity of adaptive immune regulation. For example, in a model of intestinal inflammation investigators showed that NE and a β1 agonist significantly suppressed IFN-γ and TNF-α production from the intestinal lymphocytes, with a β1 antagonist reversing these effects (32). While this study still demonstrates an immunosuppressive role for adrenergic signaling, it provided early evidence that not all impacts were orchestrated via an exclusive β2 receptor-mediated pathway. Additionally, early studies suggested that alpha receptors were not expressed on T-lymphocytes, likely due to experimental designs using alpha agonists that did not show significant changes in T-lymphocyte activation (33). However, this assumption has since been challenged with current research showing that both alpha 1 (α1) and 2 (α1) receptors are in fact expressed on T-lymphocytes from the thymus, spleen, and peripheral blood, further suggesting that their functional role is likely context-dependent (34, 35). The question still remains though as to what is the specific context that determines the impact of adrenergic receptor signaling on T-lymphocytes?

Our working hypothesis to this complex question possibly lies in cAMP. Both β1 and β2 adrenergic receptors signal through Gs proteins, which trigger adenylate cyclase to generate cAMP. In contrast, α1 receptors signal Gq proteins, which primarily signal via calcium. During classical antigen presentation, calcium signaling is a primary mediator of T-lymphocyte activation and is in incredibly high abundance. Therefore, if α1 receptors were stimulated at the same time as antigen presentation, the effect of the adrenergic signaling may be overshadowed by the already plentiful calcium signaling. This may explain why many studies have demonstrated minimal effects of alpha adrenergic signaling on T-lymphocytes, and why it is often discounted as not playing a critical role in the modulation of adaptive immune inflammation. In contrast, cAMP is not a primary component of classical signaling via the T-lymphocyte receptor, and could significantly alter the intracellular milieu of activated second messenger proteins, especially if there are no other signals present at the time of activation (i.e., cytokine receptor signaling). Moreover, since both β1 and β2 receptors signal via cAMP, they may act as redundant backups on T-lymphocytes, whereas loss of one is compensated by the other. This is supported by our previous report demonstrating that combined β1/2 antagonism was necessary to limit IL-17A levels from T-lymphocytes and that selective β1 or β2 blockade had no impact (15). This is further reinforced by the abundance of studies demonstrating other cAMP related receptors, such as dopamine or serotonin receptors, on T-lymphocytes play significant roles in shaping their polarization and inflammatory capacities, including IL-17A production (1, 36–38). While appearing unique mechanisms when examined individually, when taken together, these studies all suggest that cAMP acts to fine tune T-lymphocyte polarization and inflammation independent of the specific receptor that increases/decreases its abundance. However, the specific output of how cAMP modulates T-lymphocytes also appears highly dependent upon the additional stimulation the cell is receiving at the time. For example, in reductionist conditions where only the T-lymphocyte receptor is engaged, cAMP increases appear to be anti-inflammatory. In contrast, if the T-lymphocyte receptor is engaged simultaneously with numerous other cytokine receptors (e.g., T_H_17 polarization conditions), the cAMP levels appear to promote a more pro-inflammatory phenotype. These conditional nuances are critical as we advance our understanding of neurotransmitter modulation of adaptive immunity and as we attempt to translate our findings into therapeutic interventions.

However, even though the autonomic regulation of T-lymphocytes is highly dynamic and complex, the study herein built off our previous discovery that combined β1/2 antagonism significantly hampers T-lymphocyte IL-17A, while promoting anti-inflammatory IL-10, production (15). Here, we aimed to investigate the role of β1/2 antagonism as a therapeutic utility in autoimmune diseases such as MS, given that the balance between pro-inflammatory T_H_17 cells and immunosuppressive Treg cells plays a pivotal role in the pathophysiology of this disease. Our findings presented align with our previous work demonstrating that β1/2 blockade can alter this balance, but more importantly, antagonism of β1/2 signaling significantly reduced disease severity. These results suggest that adrenergic signaling influences both the inflammatory and regulatory aspects of T-lymphocyte responses in autoimmunity while also elucidating β1/2 antagonism as a promising clinical utility for the management of these diseases.

Previous research into the regulatory role of the nervous system in immune responses, particularly in autoimmune diseases like experimental autoimmune encephalomyelitis (EAE), has established that the sympathetic arm of the nervous system plays a crucial role. Early studies, such as those examining the effects of chemical sympathectomy using 6-hydroxydopamine (6-OHDA), found that complete removal of sympathetic innervation led to increased disease severity in EAE (39, 40). These findings initially suggested targeting the sympathetic nervous system may have a negative impact on disease outcomes. However, there are important confounders to consider when interpreting these early studies. Specifically, these studies did not account for the autonomic control of inflammation through the cholinergic anti-inflammatory response, a mechanism that had not been well understood or thoroughly investigated at the time. As previously discussed, the cholinergic anti-inflammatory response involves catecholamine signaling on a specific subset of T-lymphocytes that produce acetylcholine, which plays a crucial role in limiting inflammation (9, 41, 42). Therefore, by eliminating sympathetic signaling through chemical sympathectomy, these early studies likely inadvertently disrupted the cholinergic anti-inflammatory pathway, limiting the ability of T-lymphocytes to produce anti-inflammatory signals. As a result, the absence of sympathetic control in these studies likely exacerbated disease outcomes, which convoluted the more nuanced role of the sympathetic nervous system in regulating EAE.

With this, investigations into the role of β adrenergic receptors in EAE have also been explored, but with mixed results. For example, early work using β1/2 agonists demonstrated a significant suppression of EAE progression (43). At face value, this finding would appear to be in direct contrast to our work presented herein, but there is an interesting caveat at play. The authors of this work clearly state that β1/2 agonism had a more pronounced effect when given either from the time of immunization or at approximately 14 days post-immunization. Moreover, when given within the first 7 days after immunization, β1/2 agonism appeared to have a minimal impact on EAE outcomes. In contrast to our work, β1/2 antagonism appeared to have a greater impact when given approximately 7 days post-immunization (also supported by another recent study in rats using a similar timeline (44)), and treatment with β1/2 antagonists at the time of immunization showed minimal influence on EAE disease progression (data not shown). This is further supported by the whole body β1/2^-/-^ mice appearing to show worse disease progression than those treated with β1/2 antagonists at 7 days post immunization. Altogether, these data support that timing appears critical in how adrenergic signaling impacts the progression of EAE, with early β1/2 agonism likely suppresses innate immune cells whereas later β1/2 antagonism limits adaptive immune cell inflammation.

Current therapies for multiple sclerosis (MS) often target T-lymphocytes, either by depleting these immune cells directly, interfering with specific molecular pathways that regulate their activity, or by limiting their effector cytokines. For example, therapies such as alemtuzumab and natalizumab are designed to modulate T-lymphocyte function. Alemtuzumab works by targeting CD52, leading to the depletion of immune cells, while natalizumab inhibits the α4-integrin subunit, preventing T-cells from crossing the blood-brain barrier (45). These therapies, while effective in managing MS symptoms, are not without their drawbacks. Specifically, natalizumab has been associated with the accumulation of T_H_17 cells expressing the melanoma cell adhesion molecule in the cerebrospinal fluid (CSF) of patients, which appears to be an adaptation to bypass the natalizumab inhibition of α4-integrin subunit and continue to traffic to the central nervous system (46, 47). Additionally, about half of the patients experience significant clinical deterioration after discontinuing natalizumab therapy (48), which underscores the need for better alternatives or adjuvants to maintain disease control.

Newer therapies are being developed to address these issues. One such therapy is secukinumab, a monoclonal antibody that neutralizes IL-17A. Secukinumab has already been approved for use in psoriasis (49) and rheumatoid arthritis (50), and it was initially being tested in MS. However, clinical trials in MS were terminated early due to the development of a more effective antibody, called CJM112, with superior potential to secukinumab. A phase II study entitled Monoclonal AntiBody and fINGOlimod (MABINGO) was planned to test CJM112 in 2015, but no data appear available as to its outcome to date (51). Despite its potential, there are concerns about the use of secukinumab, as clinical studies have shown an increased risk of suicidal thoughts and completed suicides in patients undergoing treatment (45). Thus, while therapies targeting IL-17A show promise, their side effects remain a possible concern.

In this context, our proposed therapy involving β1/2 adrenergic antagonism could offer a novel and safer alternative. Based on our previous findings, we know that β1/2 adrenergic antagonism shifts the balance between T_reg_ cells and T_H_17 cells, favoring the generation of T_regs_, which are essential for maintaining immune tolerance and regulating inflammation. The promotion of T_reg_ cells over T_H_17 cells could mitigate the pathogenic autoimmune response in MS and other T-lymphocyte-driven diseases. Additionally, one of the key advantages of drugs like propranolol is that they are already FDA-approved with well-established safety profiles. These beta blockers are already used in a variety of clinical settings, including hypertension and anxiety disorders (52–54). This makes it an attractive candidate for repurposing in MS treatment, as it could potentially be combined with existing therapies targeting IL-17A or other T-lymphocyte pathways. By using propranolol as an adjuvant treatment, we could enhance the efficacy of antibody therapies that target IL-17A, like secukinumab, while simultaneously promoting T_regs_ and reducing the harmful effects of T_H_17 cells. This combination approach could lead to a more personalized and effective treatment strategy, offering a promising avenue for improving MS therapy and potentially other autoimmune diseases driven by T-lymphocytes.

In conclusion, our study underscores the critical role of β1/2-adrenergic signaling specifically in the regulation of the T_H_17/T_reg_ balance and its potential as a therapeutic target in autoimmune diseases like MS. By modulating the balance between pro-inflammatory T_H_17 cells and anti-inflammatory T_reg_ cells, non-specific β-adrenergic blockers may offer a novel approach to managing immune-driven disorders either alone or in combination with current therapeutic approaches. However, additional research is needed to fully characterize the mechanisms underlying adrenergic receptor signaling in immune responses, particularly in relation to receptor subtype specificity, disease stage, and the interplay with other immune cells. As our understanding of the neuroimmune interface continues to evolve, β-adrenergic modulation may emerge as a promising strategy for treating autoimmune and inflammatory diseases.

## Author contributions

THL and AJC designed research studies. All authors conducted experiments, acquired data, and/or performed analyses. THL and AJC wrote the manuscript, while all authors approved the final version of the manuscript. AJC provided funding and experimental oversight.

